# Simulation of spontaneous G protein activation reveals a new intermediate driving GDP unbinding

**DOI:** 10.1101/306647

**Authors:** Xianqiang Sun, Sukrit Singh, Kendall J. Blumer, Gregory R. Bowman

## Abstract

Activation of heterotrimeric G proteins is a key step in many signaling cascades. However, a complete mechanism for this process, which requires allosteric communication between binding sites that are ~30 Å apart, remains elusive. We construct an atomically-detailed model of G protein activation by combining three powerful computational methods; metadynamics, Markov state models (MSMs), and CARDS analysis of correlated motions. We uncover a mechanism that is consistent with a wide variety of structural and biochemical data. Surprisingly, the rate-limiting step for GDP release correlates with tilting rather than translation of the GPCR-binding helix 5. β-Strands 1-3 and helix 1 emerge as hubs in the allosteric network that links conformational changes in the GPCR-binding site to disordering of the distal nucleotide-binding site and consequent GDP release. Our approach and insights provide foundations for understanding disease-implicated G protein mutants, illuminating slow events in allosteric networks, and examining unbinding processes with slow off-rates.

---

Heterotrimeric G proteins are molecular switches that play pivotal roles in signaling processes from vision to olfaction and neurotransmission.^1–3^ By default, a G protein adopts an inactive state in which guanosine diphosphate (GDP) binds between the Ras-like and helical domains of the α-subunit (Gα, Fig. 1). A dimer consisting of the β- and γ-subunits (Gβγ) also binds Gα. G protein-coupled receptors (GPCRs) trigger G protein activation by binding Gα and stimulating GDP release, followed by GTP binding to Gα and dissociation of Gα from Gβγ. Gα and Gβγ then interact with effectors that trigger downstream cellular responses. Gα returns to the inactive state by hydrolyzing GTP to GDP and rebinding Gβγ. Given the central role Gα plays, a common Gα numbering scheme (CGN) has been established to facilitate discussion of the activation mechanisms of different Gα homologs.^4^ For example, the notation Lys52^G.H1.1^ indicates that Lys52 is the first residue in helix 1 (H1) of the Ras-like domain (also called the GTPase domain, or G). S6 refers to β-strand 6 and s6h5 refers to the loop between S6 and H5.

**Figure 1.**
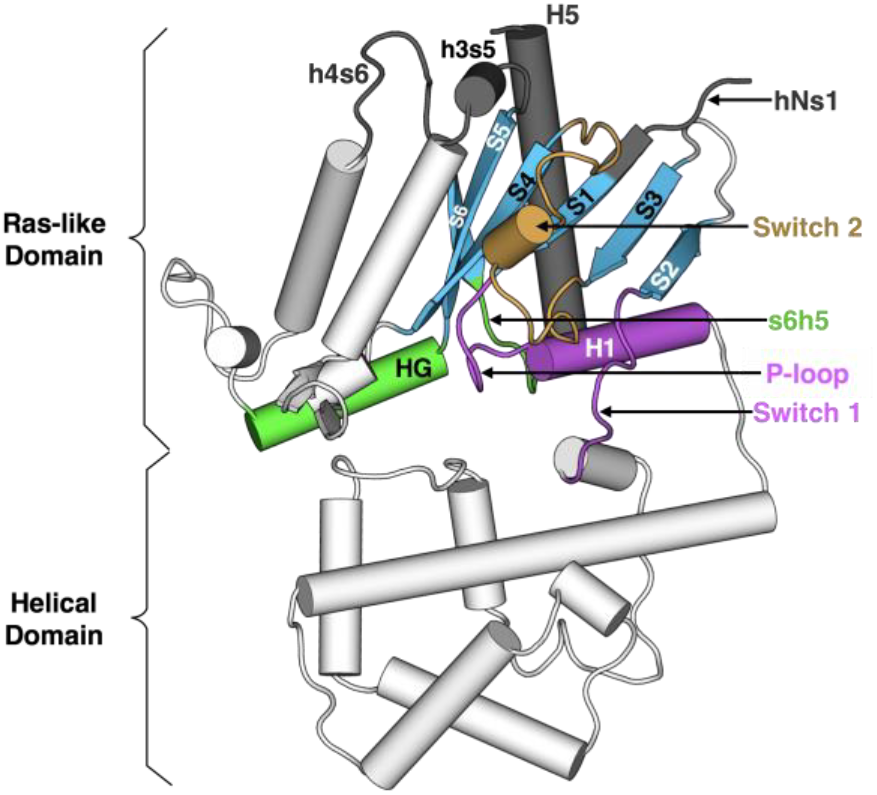
Structure of Gαq with key secondary structure elements labeled according to the Common Gα Numbering (CGN) system. The coloring scheme highlights the GPCR binding interface (gray), GDP phosphate-binding regions (pink), GDP nucleotide-binding regions (green), β-sheets (blue), and switch 2 (orange).

Strikingly, the GPCR- and nucleotide-binding sites of Gα are ~30 Å apart (Fig. 1), but the allosteric mechanism coupling these sites to evoke GDP release remains incompletely understood.^2^ Biochemical and structural studies have elucidated some key steps, but the entire process has yet to be described in atomic detail. Early studies of Gα subunits revealed structures of the GDP- and GTP-bound states, as well as the transition state for GTP hydrolysis.^5–7^ The high similarity of these structures and the binding of GDP or GTP deep in the protein's core suggests that activation occurs by adoption of other conformational states that allow GDP release.^8^ One intermediate in G protein activation was suggested by the first crystal structure of a GPCR-bound G protein in which the Ras-like and helical domains of Gα are hinged apart and GDP has dissociated.^9^ Structural analysis has led to the proposal of a universal mechanism for G protein activation.^4^ In this model, GPCR binding induces translation of H5 away from H1, which increases disorder in H1 and the P-loop (or Walker A motif^6^) to facilitate GDP release. However, there is evidence that additional intermediates may be involved in Gα activation.^2,7,10,11^ Furthermore, mutagenesis and nuclear magnetic resonance (NMR) studies have suggested important roles for other structural elements.^12–14^

Molecular dynamics simulations promise to capture the entire mechanism of G protein activation and synthesize the wealth of experimental data on this process. Methodological advances now enable simulations to capture millisecond timescale processes for proteins with less than 100 residues.^15^ For example, it is now possible to capture the binding or release of small molecules^16–20^ and peptides^21,22^ from small proteins. Such methods have simulated some of the conformational dynamics of G proteins in their inactive and active states.^23,24^ However, computational modeling of the entire process of G protein activation remains a daunting objective because the Gα subunit alone has over 300 residues and the slow kinetics of GDP dissociation (~10^−3^ min^−1^)^25^ has precluded analysis even by proprietary supercomputers.^26,27^

Here, we introduce an approach to capture infrequent or long-timescale events, such as GDP release, and reveal the mechanism of Gα activation. As a test of this methodology, we apply it to Gαq, which has one of the slowest GDP release rates^25^ and is frequently mutated in uveal melanoma.^28,29^ First, we combine two powerful sampling methods, metadynamics^30^ and Markov state models (MSMs),^31^ to observe GDP release and identify the rate-limiting step for this slow process. Then we use our recently developed CARDS method,^32^ which quantifies correlations between both the structure and disorder of different regions of a protein, to identify the allosteric network connecting the GPCR- and nucleotide-binding sites. The resulting model is consistent with a wealth of experimental data and leads to a number of predictions, described below. Taken together, our results provide the most comprehensive model of G protein activation to date. Importantly, our model was built using commodity hardware, so others can employ the methodology introduced here to study other slow events, including conformational changes and unbinding processes.

## RESULTS AND DISCUSSION

### Capturing G protein activation and GDP release in atomic detail

We reasoned that studying the mechanism of spontaneous GDP release from a truncated form of Gαq would be representative of the mechanism of GPCR-mediated activation of the heterotrimeric G protein while minimizing the computational cost of our simulations. This hypothesis was inspired by previous work demonstrating that a protein's spontaneous fluctuations are representative of the conformational changes required for the protein to perform its function.^33,34^ Therefore, we hypothesized that GPCRs stabilize conformational states that G proteins naturally, albeit infrequently, adopt in the absence of a receptor. We chose to focus on Gα since it forms substantial interactions with GPCRs, compared to the relatively negligible interactions between GPCRs and G protein β and y subunits. This view is supported by the fact that GPCRs and ‘mini’ G proteins, essentially composed of just the Ras-like domain of Gα, recapitulate many features of GPCR-G protein interactions.^35^ We also reasoned that truncating the last 5 residues of Gαq would facilitate the activation process because these residues contact Gα in some GDP-bound structures,^36,37^ contact the GPCR but not the G protein in the GPCR-bound state, and removing them promotes GDP release due to a reduced GDP-binding affinity.^38,39^ Taken together, such evidence suggests that the C-terminus interacts with the rest of of Gα to stabilize the inactive state and that these interactions are disrupted by binding of an activated GPCR. In support of this hypothesis, we find that the energetic barrier to GDP release is lower in metadynamics simulations of the truncated variant than for full-length Gαq (Fig. S1). These simulations, and those described hereafter, were initiated from an x-ray structure of the Gαq heterotrimer bound to GDP and an inhibitor of nucleotide exchange^40^; Gβγ and the inhibitor were excluded from all simulations.

To observe G protein activation, we developed a variant of adaptive seeding^41^ capable of capturing slow processes like ligand unbinding that are beyond reach of conventional simulation methods. First, we use metadynamics^20,30,42^ to facilitate broad sampling of conformational space by biasing simulations to sample conformations with different distances between GDP and Gαq. Doing so forces GDP release to occur but provides limited mechanistic information because adding a biasing force can distort the system's kinetics or cause the simulations to sample high-energy conformations that are not representative of behavior at thermal equilibrium. To overcome these limitations, we chose starting conformations along release pathways observed by metadynamics as starting points for standard molecular dynamics simulations, together yielding an aggregate simulation time of inn.6 µs. These simulations should quickly relax away from high-energy conformations and provide more accurate kinetics. Then we use these simulations to build an MSM. MSMs are adept at extracting both thermodynamic *and* kinetic information from many standard simulations that, together, cover larger regions of conformational space than any individual simulation.^31^ Related approaches have successfully captured the dynamics of small model systems^43^ and RNA polymerase.^44^

This protocol enabled us to capture the entire mechanism of G protein activation, including GDP release and the rate-limiting step for this process, without requiring specialized hardware. Identifying the rate-limiting step for this process is of great value because GDP release is the rate-limiting step for G protein activation and downstream signaling. Therefore, the key structural and dynamical changes responsible for activation should be apparent from the rate-limiting conformational transition for this dissociation process.

To identify the rate-limiting step, we applied transition path theory^45,46^ to find the highest flux paths from bound structures resembling the GDP-bound crystal structure to fully dissociated conformations. Then we identified the least probable steps along the 10 highest flux release pathways (Fig. 2A and S2), which represent the rate-limiting step of release. By comparing the structures before and after the rate-limiting step, we define the bound state as all conformations where the distance from the center of mass of GDP's phosphates to the center of mass of three residues on H1 that contact the GDP β-phosphate (Lys5n^G.H1.1^ \ Ser53^G.H1.2^, and Thr54^G.H.1.3^) is less than 8Å. Consistent with this definition, this distance remains less than 8Å throughout the entirety of 35.3 µs of GDP-bound simulations.

**Figure 2.**
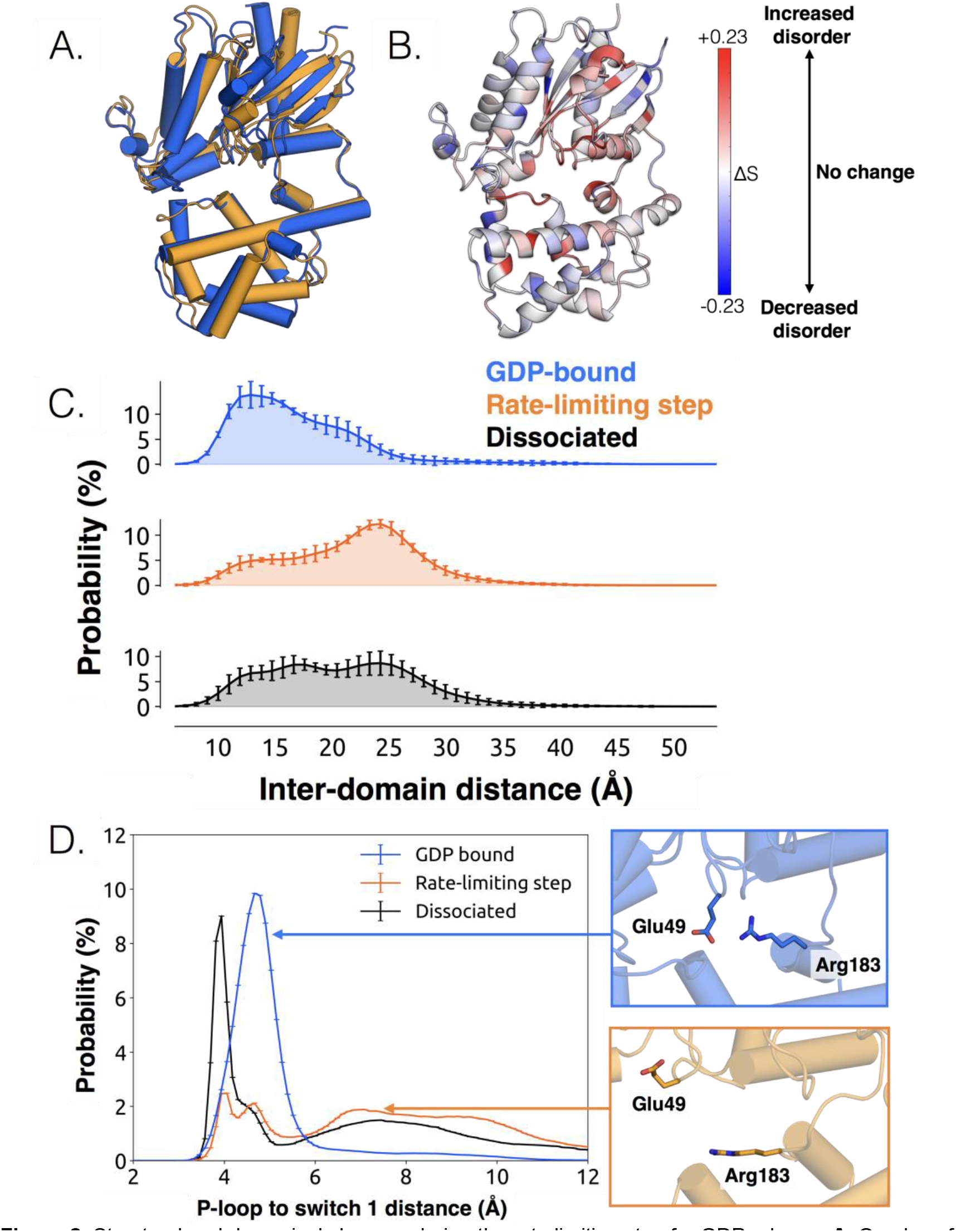
Structural and dynamical changes during the rate limiting step for GDP release. **A.** Overlay of representative structures before (blue) and after (orange) the rate limiting step. **B.** Change in conformational disorder (Shannon's entropy) across the rate-limiting step, according to the color scale on the right. **C.** Histograms of interdomain distances before (top, blue) and after (middle, orange) the rate limiting step, along with the inter-domain distance distribution after GDP is released (bottom, blacky Inter-domain distance is measured using residues analogous to those used in DEER experiments,^23^ Leu97^H.HA.29^ in the helical domain and GIu 250^G.H3.4^ on H3. **D.** Distribution of distances between Glu49^G.H3.4^ and Arg183^G.hfs2.2^ (left) before (blue) and after (orange) the rate-limiting step, as well as after GDP release (black). Representative structures of the interacting residues are also shown (right).

The conformational changes we observe during the rate-limiting step are consistent with previous structural and biochemical work. For example, we observe that the Ras-like and helical domains separate (Fig. 2C), as observed in DEER experiments^47^ and simulations.^23^ This finding is consistent with the intuition that these domains must separate to sterically permit GDP release. Domain opening is accompanied by disruption of a key salt bridge between Glu49^G.s.1h1.4^ of the P-loop and Arg183^G.hfs2.2^ of switch 1 (Fig. 2D), as well as an increase in the disorder of many of the surrounding residues (Fig. 2B, 2C and S3A), consistent with the proposal that this salt bridge stabilizes the closed, GDP-bound state.^48^

While domain opening is necessary for GDP release, previous work suggests it is insufficient for unbinding.^23^ Indeed, we also see that this opening is necessary but not sufficient for GDP unbinding, as the Ras-like and helical domains often separate prior to release (Fig. 2C). Notably, the Ras-like and helical domains only separate by ~10 Å during the rate-limiting step. In contrast, these domains separate by 56 Å in the first structure of a GPCR-G protein complex. This result suggests that GDP release may have occurred long before a G protein adopts the sort of widely opened conformations observed in crystallographic structures.^9^

### Tilting of H5 helps induce GDP release

We also observe less expected conformational changes associated with GDP release. The most striking is tilting of H5 by about 26° (Fig. 3A, and S4). We find that H5 tilting correlates strongly with the distance between GDP and Gαq along the highest flux dissociation pathways (Fig. 3B). In particular, substantial tilting of H5 is coincident with the rate-limiting step for GDP release. This tilting contrasts with x-ray structures and the universal mechanism, in which H5 is proposed to translate along its helical axis towards the GPCR, initiating the process of GDP release (Fig. 3A). During our simulations we also observe translation of H5, but it is not correlated with the rate-limiting step of GDP release (Fig. 3C and S5). Therefore, we are not merely missing an important role for translation due to insufficient sampling.

**Figure 3.**
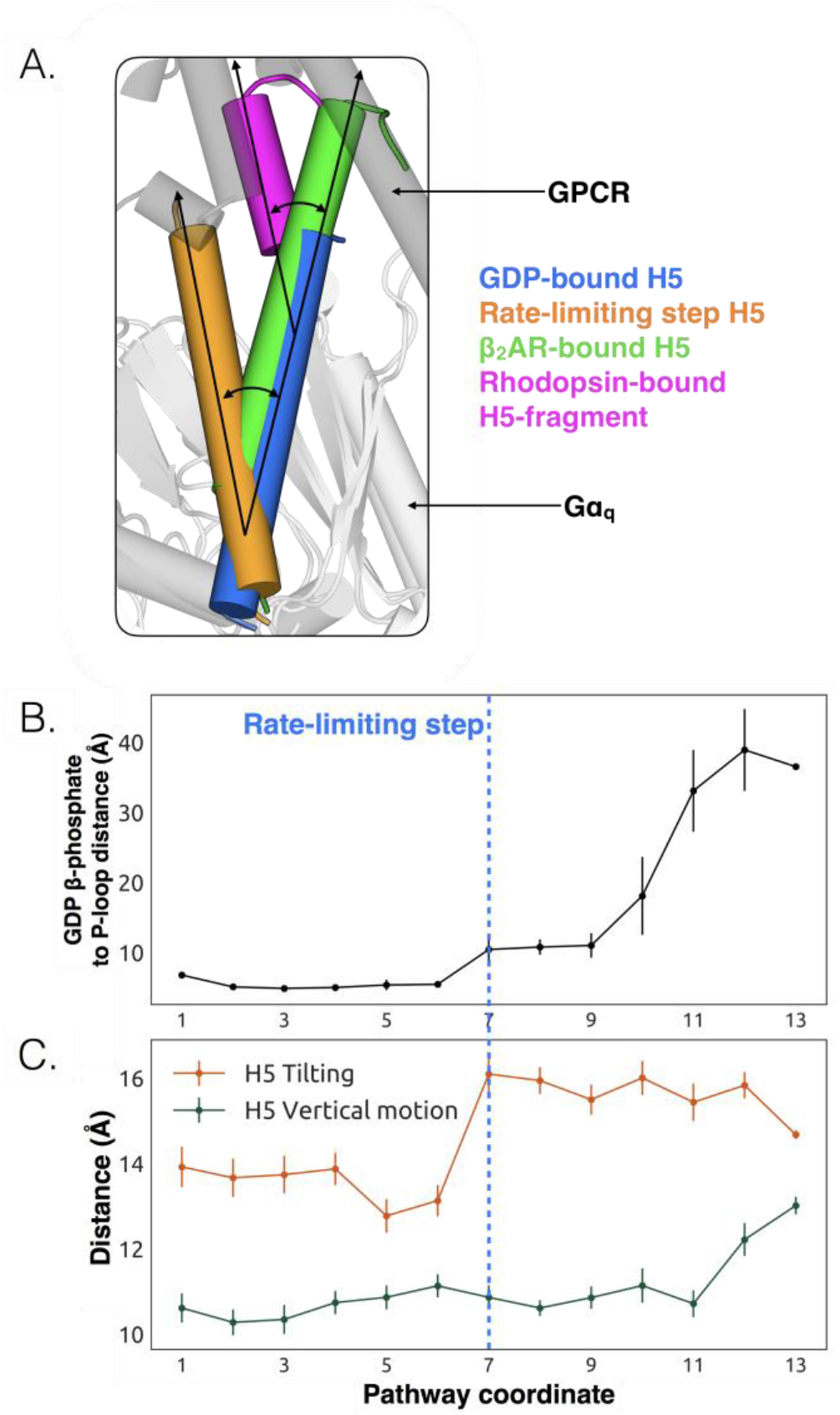
Tilting of H5 is correlated with GDP release but translation of H5 is not. **A.** Displacements of H5 relative to the GDP-bound crystal structure (blue). The three other orientations of H5 come from after the rate limiting step in our model (orange), the co-crystal structure of Gas and β2AR (green, PDB ID 3SN6)^11^, and the co-crystal structure of a C-terminal fragment of Gat and rhodopsin (magenta, PDB ID 3PQR)^30^ The black arrows highlight the change in orientation of the long axis of each helix. A representative GPCR (gray) and Gα (white) structure are shown for reference. **B.** GDP release distance across the highest flux pathway, defined as the distance from the GDP β- phosphate to the center of mass between residues Lys52^G.H1.1^, Ser53^G.H1.2^, and Thr54^G.H1.3^ on the H1. The state marking the rate-limiting step is highlighted by the blue dashed line. **C.** H5 motion across the highest flux pathway. The distances measured here representing H5 motion are taken from the same states as in B. H5 tilting (orange) is measured by the distance between Leu349^G.H5.16^ and Tyr325^G.S6.2^. Likewise, H5 vertical motion (green) is measured by the distance between Thr334^G.H5.1^ and Phe341^G.H5.8^. The state after the rate-limiting step is marked with the blue dashed line, extended down from B.

The potential importance of H5 tilting is supported by other structural data. For example, a crystal structure of rhodopsin^49^ with a C-terminal fragment from H5 of Gαt shows a similar degree of tilting (Fig. 3A). Also, the tilt of H5 varies in crystal structures of the β2AR-Gs complex,^9^ two different GLP-1 receptor-Gs complexes,^50,51^ and an A2AR-mini-Gs complex.^35^ The potential relevance of tilting has also been acknowledged by a number of recent works.^4,9,52^ Finally, H5 is translated toward the GPCR in the A2AR-mini-Gs structure but GDP remains bound. The authors of that study originally suggested that one of the mutations in mini-Gs decouples H5 translation from GDP release. However, given that we see GDP release without H5 translation in our simulations, it is also possible that amino acid substitutions required to create mini-Gs instead mitigate H5 tilting.

We propose that tilting of H5 is an early step in the GDP release process, which is followed by upward translation of this helix to form a GPCR-G protein complex primed to bind GTP. This hypothesis stems from our observation that tilting of H5 is coincident with the rate-limiting step for GDP release, while translation of H5 only becomes stable after GDP dissociates (Fig. 3C). This model is consistent with previous suggestions that G protein activation occurs through a series of sequential interactions with a GPCR.^2,9^ Another possibility is that any displacement of H5, whether tilting or translation, may be sufficient to trigger GDP release.

### Identification of the allosteric network that triggers GDP release

While conformational changes of H5 are important for Gα activation, other regions could also play a role.^11,13^ However, it is not straightforward to determine what other structural elements contribute to activation or their importance relative to H5. Our hypothesis that spontaneous motions of a protein encode functionally relevant conformational changes suggests that the coupling between the GPCR- and nucleotide-binding sites of Gα should be present in simulations of the inactive protein; This provides a means to identify key elements of this allosteric network. To test this hypothesis, we ran 35.3 microseconds of simulation of GDP-Gαq. Then we applied the CARDS method^32^ to detect correlations between both the structure and dynamical states of every pair of dihedral angles. These pairwise correlations serve as a basis for quantifying the correlation of every residue to a target site, such as the GPCR-binding site. Combining these correlations with structural and dynamical changes in our model of GDP release provides a basis for inferring how perturbations to the GPCR-binding site are transmitted to the nucleotide-binding site.

To understand how H5 tilting triggers GDP release, we first identified structural elements with strong coupling to H5 and then worked our way outward in repeated iterations until we reached the nucleotide-binding site. This analysis reveals that tilting of H5 directly communicates with and impacts the conformational preferences of the s6h5 loop, which contacts the nucleobase of GDP (Fig. S6). During the rate-limiting step, these contacts are broken and there is increased disorder in the s6h5 loop, particularly Ala331 of the TCAT motif (Fig 4 and S3B). The importance of the TCAT motif in our model is consistent with its conservation and the fact that mutating it accelerates GDP release.^53–55^ Based on our model, we propose these mutations accelerate release by weakening shape complementarity with GDP.

**Figure 4.**
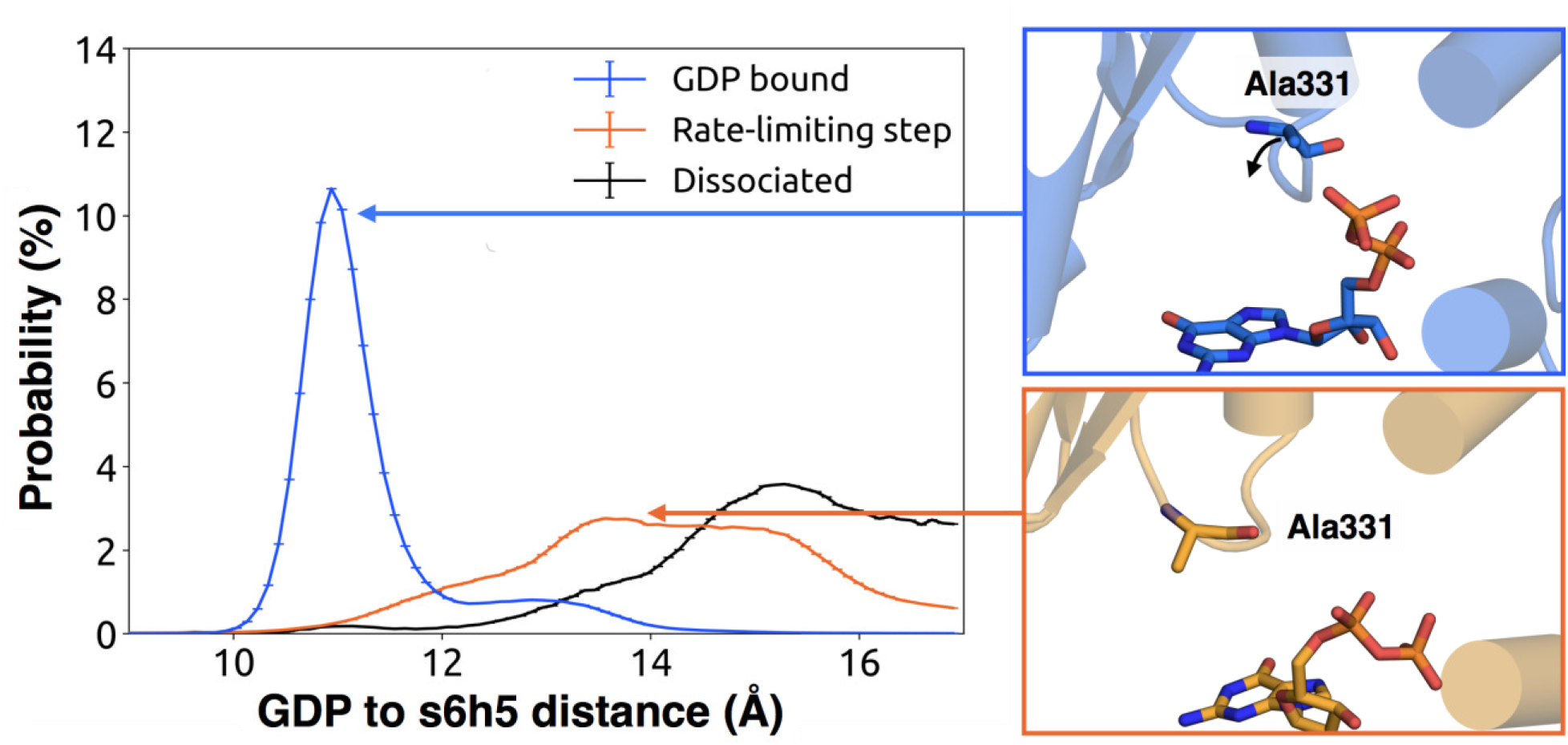
Change in the s6h5 loop conformation across the rate-limiting step. Distribution of distances (left) from GDP's β-phosphate to Ala331^G.s6h5.3^ on the s6h5 loop before (blue) and after (orange) the rate limiting step, as well as after GDP release (black). Representative structures of the s6h5 loop (right) are shown for before (top right, blue) and after (bottom right, orange) the rate limiting step.

We also observe an important role for communication from H5 to H1, consistent with the universal mechanism. In particular, H1 is strongly coupled with the s6h5 loop (Fig. S6), which is sensitive to displacement of H5. In the rate-limiting step, s6h5 moves away from H1, contributing to an increase in disorder of H1 and the P-loop (Fig. S3A and S3B). Increased disorder in a set of residues that directly contact the GDP phosphates (Glu49^G.s1h1.4^, Ser50^G.s1h1.5^, Gly51^G.s1h1.6^, Lys52^GH1.1^, and Ser53^G.H1.2^) likely contributes to a reduced affinity for GDP (Fig. S3A). Increased disorder in these residues also contributes to disruption of the salt bridge between Glu49^G.s1h1.4^ of the Ras-like domain and Arg183^G.hfs2.2^ of the helical domain, facilitating inter-domain separation.

We further note that the s6h5 loop impacts the nucleotide-binding site through allosteric coupling with the HG helix, which also contacts GDP via Lys275^G.s5hg.1^ and Asp277^G.HG.2^ (Fig S6E and S7). The disorder of both of these residues increases during the rate-limiting step, consistent with observations of increased mobility in HG upon receptor-mediated activation.^2^ There are also correlations between the P-loop and Lys275^G.s5hg.1^ on Helix G (Fig. S6E), which result from the disruption of a key salt bridge between Lys275 and Glu49^G.s1h1.4^ on the P-loop during the rate-limiting step (Fig. S7). Lys275^G.s5hg.1^ is conserved across all Gα families, suggesting it plays an important role in the stability or function of the protein. However, attempts to experimentally examine the role of this residue by mutating Lys275^G.shg.1^ have resulted in aggregation.^13^ Our simulations suggest Lys275 ^G.shg.1^ plays an important role in stabilizing the GDP-bound state and that breaking the salt bridge with Glu49^G.s1h1.4^ facilitates GDP release. This finding demonstrates the utility of our atomistic simulations, as we can examine the role of residues that are difficult to probe experimentally.

### H1 and the β-sheets are communication hubs

To identify other important structural elements in the allosteric network underlying G protein activation, we followed correlated motions emanating from other sites that are known to interact directly with GPCRs, including the hNs1 loop, the h3s5 loop, and the h4s6 loop.^2^ We find that h3s5 and h4s6 are largely isolated, suggesting they play a role in forming a stable GPCR-G protein complex but not in the allosteric mechanism that triggers GDP release. This finding is consistent with sequence analysis suggesting these structural elements are important determinants of the specificity of GPCR-Gα interactions.^56^ In contrast, the hNs1 loop has strong correlations with β-strands S1-S3 (Fig. S8). These strands, in turn, communicate with H1, switch 1, and the P-loop.

Integrating our correlation analysis with structural insight from the rate-limiting step described above suggests an important role for S1-S3 in a complex allosteric network that couples the GPCR- and nucleotide-binding sites (Fig. 5, S8, and S9). S2 and S3 twist relative to S1 and away from H1 (Fig. 2A and S10). This twisting disrupts stacking between Phe194^G.S2.6^ on S2 and His63^G.H1.12^ on H1 and increases disorder of side-chains in H1 (Fig. 2B, 6, and S3C). Increased disorder in H1 is also a crucial component of the proposed universal mechanism, but in that model translation of H5 is the key trigger for changes in H1. The role for the β-sheets in our model is consistent with NMR experiments showing chemical exchange in the methyls of S1-S3 upon receptor binding^12^ and mutational data.^57–59^

**Figure 5.**
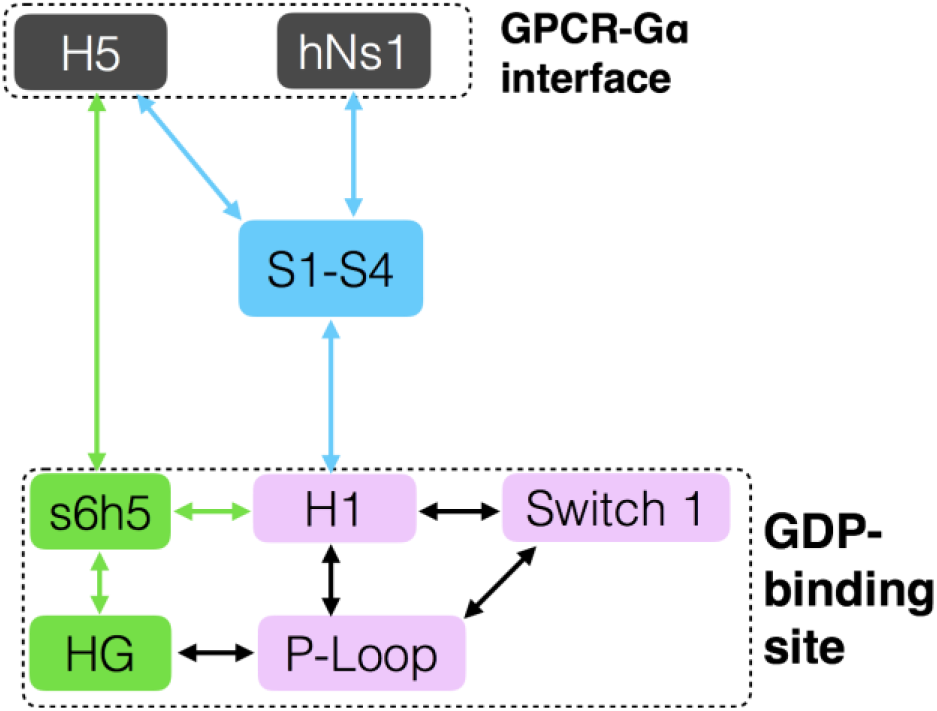
Allosteric network connecting the GPCR- and nucleotide-binding interfaces. The coloring scheme corresponds to that used in Figure 1, highlighting the GPCR binding interface (gray), GDP phosphate-binding regions (pink), GDP nucleotide-binding regions (green), and the β-sheets (blue).

**Figure 6.**
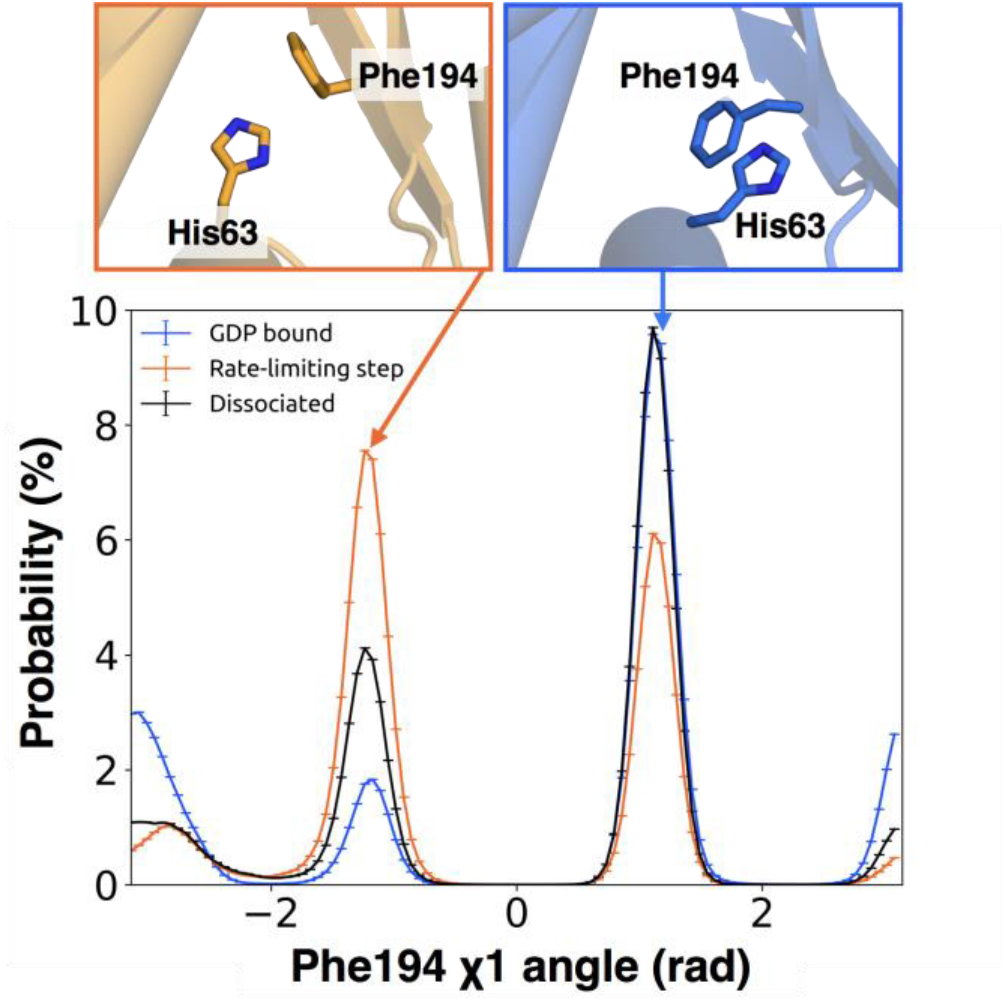
π-stacking between S2 and H1 is disrupted during the rate-limiting step. **A.** Distribution of the x1 angle (bottom) of Phe194^G.S2.6^ on S2 before (blue) and after (orange) the rate limiting step, as well as after GDP release (black). Representative structures of Phe194^G.S2.6^ and His63^G.H1.12^ (top) corresponding to before and after are also shown.

The importance of H1 and β-strands S1-S3 is underscored by mapping the global communication of every residue onto a structure of Gα (Fig. S11). The global communication of a residue is the sum of its correlations to every other residue and is a useful metric for identifying residues that are important players in allosteric networks.^32^ Interestingly, these β-strands and H1 have higher global communication than H5 and the s6h5 loop. This suggests that H1 and the β-sheets integrate conformational information from multiple sources, including the hNs1 loop, and not just H5. The importance of the β sheets and H1 for allosteric communication is consistent with their conservation,^13^ in contrast to simply promoting protein folding and stability as had been suggested by the similar conformation they adopt across static crystal structures.

### GDP release alters the structure and dynamics of the Gβ binding site

We also find that switch 2 has strong correlations with the nucleotide-binding site, especially switch 1 (Fig. S9E). Given that switch 2 is a major component of the interface between Gα and Gβ, this communication could enable GDP release to trigger dissociation of Gα from Gβγ. Examining the rate-limiting step for GDP release reveals that switch 2 shifts towards the nucleotide-binding site (Fig. 2A and S12) and exhibits increased conformational disorder (Fig. 2B and S3D). These findings are consistent with previous kinetic studies postulating that switch 2 dynamics are impacted prior to GDP release.^60^

## CONCLUSIONS

We have succeeded in simulating G protein activation, particularly the allosteric coupling between the GPCR- and nucleotide-binding sites of Gαq and consequent unbinding of GDP. Our results reveal a previously unobserved intermediate that defines the rate-limiting step for GDP release and, ultimately, G protein activation. Our model synthesizes a wealth of experimental data and previous analyses. For example, we identify an important role for coupling from H5 to the s6h5 loop and H1 that is consistent with a previously proposed universal mechanism for G protein activation. However, we also find that this allosteric network incorporates the hNs1 loop, β-strands S1-S3, and the HG helix. Strands S1-S3 and H1 serve as hubs in this network, simultaneously integrating information from both H5 and the hNs1 loop. The consistency of our model with a wide variety of structural and biochemical data suggests that it is a promising foundation for future efforts to understand the determinants of GPCR-Gα interaction specificity, how mutations cause aberrant signaling and disease, and how small molecule inhibitors modulate Gα activation. Our model also adds weight to the growing appreciation for the fact that a protein's spontaneous fluctuations encode considerable information about its functional dynamics. Construction of our model was enabled by a powerful combination of simulation methods, namely metadynamics and MSMs, using only conventional hardware. In the future, we expect this methodology will prove valuable for understanding other slow conformational changes and unbinding processes.

## ACKNOWLEDGEMENTS

We thank T.E. Frederick and T.D. Todd for their helpful discussion and insight. We are grateful to the Folding@home users for computing resources. This work was funded by National Institutes of Health grants R01GM12400701, R01GM044592 and R01GM12409301, as well as by the National Science Foundation CAREER Award MCB-1552471. G.R.B. holds a Career Award at the Scientific Interface from the Burroughs Wellcome Fund and a Packard Fellowship for Science and Engineering from The David & Lucile Packard Foundation.

## AUTHOR CONTRIBUTIONS

X.S. and S.S. contributed equally to this work. X.S., K.J.B., and G.R.B designed and ran initial simulations. X.S., S.S., K.J.B., and G.R.B analyzed the data and generated figures. X.S., S.S., K.J.B. and G.R.B. wrote the manuscript.

## METHODS

### Molecular dynamics simulations

Simulations of Gαq were initiated from PDB ID: 3AH8^40^ after removing the first 36 residues, which come from Gai. The protein was solvated with TIP3P explicit solvent in a dodecahedron box extending 1nm beyond the protein in every dimension and neutralized with Na^+^ and Cl^−^. A single Mg^2+^ ion was added coordinating the phosphate groups of GDP as its presence accelerates GDP release.^61^ Simulations were run with Gromacs^62^ using the AMBER03^63^ Force field and GDP parameters obtained from the AMBER Parameter Database^64^ (http://research.bmh.manchester.ac.uk/bryce/amber). Details of the minimization and equilibration steps are provided in the SI. Production runs were carried out at a temperature of 300K and a pressure of 1 bar using the v-rescale thermostat^65^ and the Parrinello-Rahman barostat.^66^ Virtual sites^67^ were used to allow for a 4fs time-step.

We used a three-step protocol to simulate GDP release. First, we performed 100 parallel simulations of the GDP-bound state for an aggregate simulation time of 35.3 µs. We then selected three representative conformations at different interdomain-opening distances as starting conformations for metadynamics^30^ simulations using PLUMED.^68^ We defined two collective variables for our metadynamics simulations: 1) the distance between GDP's phosphate groups and the backbone of Lys52-Thr54 in Gαq and 2) the RMSD of GDP to the starting conformation. Additional details for our metadynamics simulations are provided in the SI and Table S1. We extracted 2,085 conformations along potential GDP release pathways identified from our metadynamics simulations using the string method. ^9^ Finally, we used these conformations as starting points for unbiased MD simulations run on Folding@home^70^ for an aggregate simulation time of 122.6 µs.

### Identifying the allosteric network with CARDS

We applied the CARDS methodology^32^ to simulations of the GDP-bound state of Gαq. CARDS measures communication between every pair of dihedrals via correlated changes in structural motions and dynamical behavior. Structural states are captured by discretizing backbone *ϕ* and *ψ* dihedrals into two states (cis and trans) while sidechain *χ* angles are divided into three states (gauche+, gauche-, and trans). Every dihedral is also parsed into dynamical states, capturing whether the dihedral is stable in a single state (ordered), or rapidly transitioning between multiple states (disordered). Then the holistic communication is computed to quantify the correlation between the structural and dynamical states of every pair of dihedrals (SI methods).

From the pairwise correlation for every dihedral-pair, we extracted how much each individual residue communicates with a target site of interest. After locating the group of residues communicating most strongly with a specific target site, we set this newly identified group as the new target site; The iteration of this process allows us to identify a pathway of communication from one region of interest to another. Here, we set the GPCR contact sites as our initial target sites. We then iteratively used this approach to identify pathways connecting these contact sites with the GDP-binding site of Gαq.

### Markov state model construction

We clustered Gαq conformations and GDP binding states separately and combined the assignments to build a Markov State Model using MSMBuilder (v2.8).^71,72^ First, we clustered protein conformations into 5040 states using a hybrid k-centers/k-medoids method with a 1.8 Å cutoff. Then we clustered the GDP-binding state into 321 states using the automatic partitioning algorithm (APM)^73^ with a residence time of 2ns. By combining the assignments from protein conformations and the GDP binding state, we obtained a total of 221,965 states. The implied timescales of this MSM show Markovian behavior with a lag time of 5ns (Figure S13).

### Quantifying conformational disorder

The disorder of every residue was measured by computing Shannon entropy^74^ of each dihedral as they are natural degrees of freedom for describing protein dynamics. Shannon entropy (*H*) is defined as

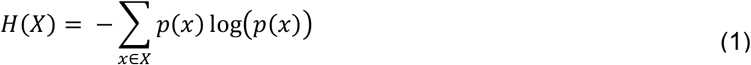

where *x* ϵ *X*refers to the set of possible states that dihedral *X* can adopt and *p*(*x*) is the probability that dihedral *X* adopts state *x*. Dihedral angles were calculated using MDTraj^75^ and assigned to discrete rotameric states as described above using CARDS. The entropy of a single residue was computed by summing up the entropies of its dihedrals, and normalized by the residue's maximum possible Shannon entropy.

### Identification of the rate-limiting step for GDP release

We used transition pathway theory (TPT)^45,46^ to find the highest flux paths from the bound state to the unbound state. The bound state was defined as all clusters that satisfied two criteria: (i) GDP is within 6 Å of the backbone atoms of Lys52-Thr54 and (ii) GDP has an RMSD <0.5 Å to its crystallographic conformation. The unbound state was defined as all clusters with GDP >55 Å from the binding pocket. The rate-limiting step was identified by finding the bottlenecks of the highest flux paths.

